# DiffTAD: Detecting Differential contact frequency in Topologically Associating Domains Hi-C experiments between conditions

**DOI:** 10.1101/093625

**Authors:** Rafal Zaborowski, Bartek Wilczynski

## Abstract

**Motivation:** In recent years, the interest in analyzing chromosome conformation by Hi-C and related techniques has grown. It has been shown that contact frequency matrices obtained by these methods correlate with other methods of measurement of activity such as transcriptomics and histone modification assays. This brings a question of testing for differential contact frequency between experiments to the field.

**Results:** In this work, we provide a freely available software that implements two statistical methods for testing the significance of differential contact frequency in topological domains between two experiments. One method follows an empirical, permutation based approach to computing p-values, while the other is a parametric test based on the Poisson-Binomial distribution.

**Availability:** The software is freely available on the GNU General Public License at https://bitbucket.org/rzaborowski/differential-analysis

**Contact:** [r.zaborowski|bartek]@mimuw.edu.pl

**Supplementary information:** Supplementary data are available at *Bioinformatics* online.

## 1 Introduction

Hi-C is a method to study global chromosomal architecture by analysis of chromatin contacts ([Lieberman-Aiden *et al*., 2009]). As a result of Hi-C assay a library of paired end reads is produced. Reads are then mapped onto reference genome and interpreted as pairs of spatially neighbouring regions. Hi-C data is usually analyzed with help of contact maps - symmetric matrices where rows and columns mark chromosomal regions and cells on their intersection represents number of contacts between them. A particular feature associated with contact maps of eukaryotic genomes is presence of Topologically Associating Domains (TADs) - regions of genome strongly enriched with self-interactions ([Dixon *et al*., 2012]). The role of TADs is still not fully understood however it has been shown that some TADs strongly correlate with replication domains ([Pope *et al*., 2014]) and separate different regulatory neighbourhoods ([Andrey *et al*., 2013]). The deletion of TAD boundaries can lead to disruption of large chromosomal landscapes and consequently cause transcriptional misregulation ([Nora *et al.*, 2012]). Moreover TAD boundaries tend to show significant level of conservancy between cell types and species ([Dixon *et al.*, 2012]).

Recently Hi-C became a useful technique for differential analysis. This sort of analysis aim to explain gene expression or epigenetic modification differences between cell types, species or in response to experimental conditions like different stimuli treatment. For example in [Le Dily *et al.*, 2014] and [Dixon *et al.*, 2015] it was shown that depletion and enrichment in Hi-C intra-TAD interactions is linked with genes down- or upregulation respectively. This kind of studies require appropriate statistical methods for detection and quantification of chromatin regions with depletion or enrichment in interactions.

In this paper we present an application for systematic detection of TADs enriched or depleted in contact frequency. Our pipeline implements two methods. First is based on permutation test and is very similar to the one used in [Dixon *et al.*, 2015]. Second method uses parametric test based on Poisson Binomial distribution.

## 2 Definitions

In this article we will use *E* = {*A, B*} to represent a pair of different experiments *A* and *B*. This can be different cell lines or different experimental conditions for example treatment with stimuli and no treatment, etc. Each experiment *L* ∈ *E* is a tuple: *L* = (*M^L^*, *T^L^*), where 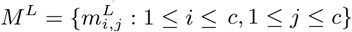, *c* represents chromosome length in given resolution and *m^L^* is symmetric *c* × *c* matrix such that: 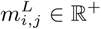. For simplicity we present a case for one chromosome. In case of many chromosomes the reasoning is analogous. *T^L^* = {*t^L^*: (*i, s, e*), *i* ∈ ℕ, 1 ≤ *s* ≤ *c*, 1 ≤ *e* ≤ *c*}, where *i* is TAD identifier and *s*, *e* are TAD start and TAD end respectively. In other words each experiment consists of set of contact maps and set of TADs.

## 3 Methods

### 3.1 Preprocessing

Pipeline input are 2 sets of experimental data *A* and *B*. First step of preprocessing is to calculate a sum of contacts for each map of experiments under study: 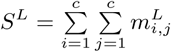. Next each contact map is normalized by dividing all its entries by appropriate sum and multiplying by constant factor: 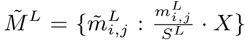, where constant factor *X* is equal to the mean of *A* and *B* contact sums, i.e.: 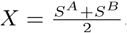. Lastly differential map is computed according to formula: 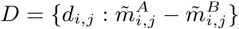.

### 3.2 Permutation test

In this approach the type of domain and strength of effect (enrichment or depletion) is calculated using the median of TAD differential interactions. Using such test statistic positive median represent enriched TAD with respect to experiment *A* and negative median means that TAD is enriched in experiment *B* relative to *A*. To assess strength of enrichment or depletion of TADs a distribution of medians in randomized data for each TAD is produced:

1. for each chromosome calculate *N* random differential maps by randomly permuting all entries in the matrix along their respective diagonals,
2. based on random differential maps calculate distribution of medians for each TAD.

Given distribution of randomized medians for each TAD we can calculate an empirical p-value of observing actual TAD median differential interactions. Finally obtained p-values are corrected for multiple hypothesis testing using Bonfferoni correction.

### 3.3 Parametric test

Another method for differential domains detection can be based on the following intuition. If a TAD is enriched in contacts relative to experiment *A* we would see most of its cells positive. We can associate this pattern with flipping a coin *N* times with probabilities *p^TAD^* = (*p*_1_, …, *p_N_*) and observing how often (*n*) we see heads (positive values). In this case 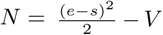 refer to the number of cells inside TAD, 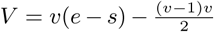and *v* stands for the number of diagonals excluded from contact map starting from main diagonal (i.e. masked diagonals). Due to Hi-C contact decay bias we estimate vector of probabilities for each differential map *p^D^* = (*p*_*v*+1_, …, *p_c_*) by calculating *p_i_* for the *i*-th diagonal separately. *p_i_* is then calculated by dividing number of positive (negative for depletion) bins on *i*-th diagonal by the number of all bins on this diagonal. Our test statistic for a TAD of size *N* is then number of positive bins (negative for testing depleted TADs) and we calculate p-value of observing this number of positive bins assuming Poisson-Binomial distribution. Due to the fact that 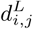 do not have binary values, but reals we approximate *n* using following formula:

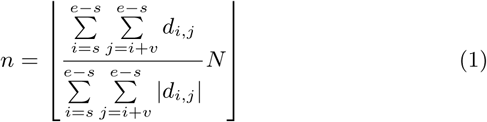

The rationale here is that higher the values of cells inside TAD, more enriched it gets. Conversely if TAD contains negative cells it lowers its enrichment effect and therefore lowers *n*. Assuming that *n* ∼ *PB*(*k, N, p^TAD^*) we want to calculate: *Pr_PB_* (*k* ≥ *k,N,p^TAD^*). Calculated p-values are then corrected for multiple hypothesis testing using Bonferroni correction.

## 4 Sample usage

Below we present an example of our method applied to differential map of human chromosome 1 (fig. 1). Upper triangle part illustrates comparison between mesenchymal stem cells (MSC) and embryonic stem cells (ESC) with detected enriched and depleted TADs. For comparison lower part presents MSC replicates difference. The number of detected TADs is equal to 61 enriched and 73 depleted in first case while MSC replicates comparison produced 43 enriched and 20 depleted TADs. Hi-C data was taken from [Dixon *et al.*, 2015]. TADs used for analysis were produced using Armatus software ([Filippova *et al.*, 2014]) applied to MSC contact maps. The test used was parametric with p-value threshold 0.05.

**Figure 1:**
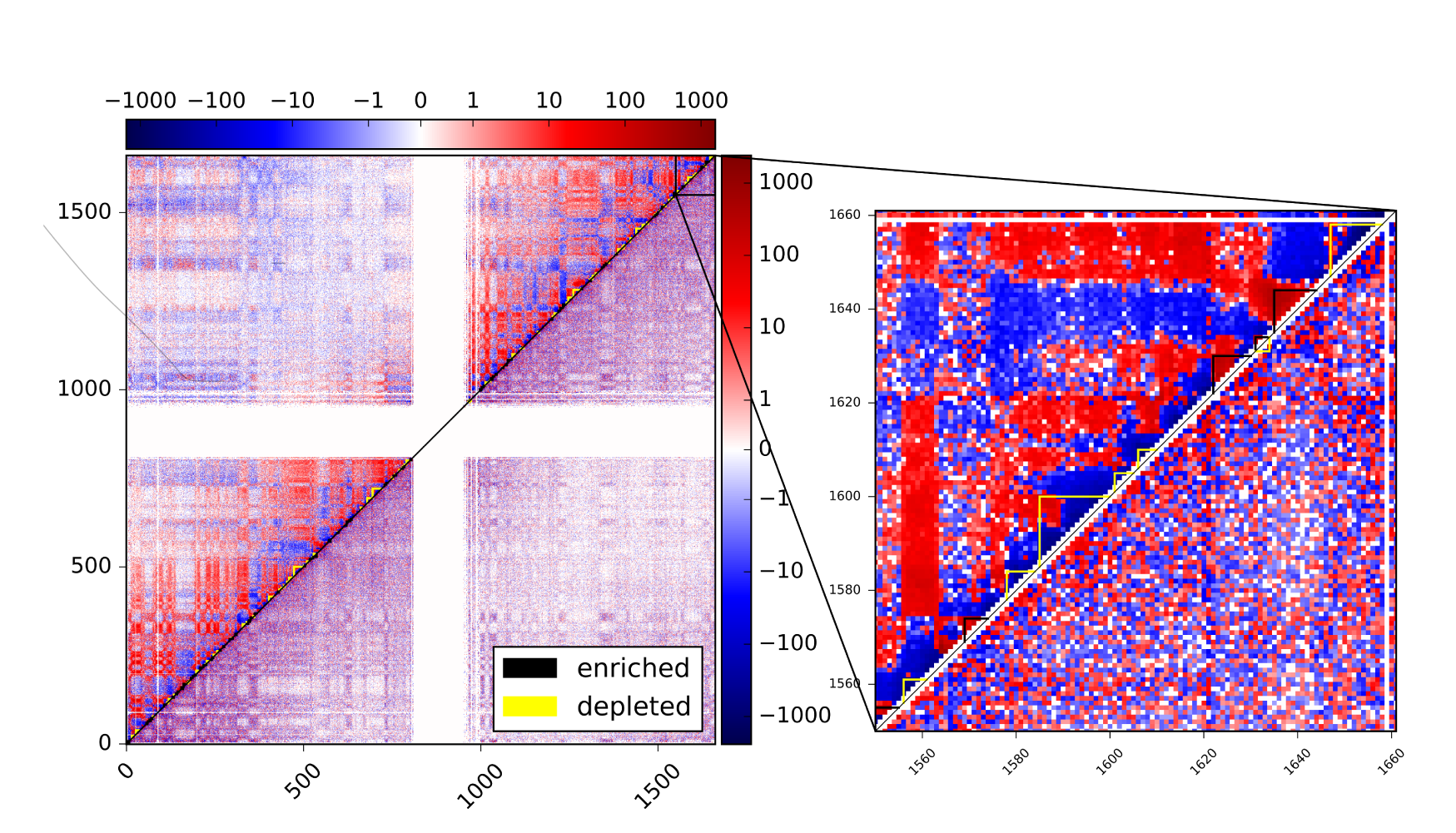
Differential map of human chromosome 1. Above diagonal: MSC replicate 1 - ESC replicate 1, below diagonal: MSC replicate 1 - MSC replicate 2.

## 5 Discussion and Conclusion

The software we present here allows a growing group of researchers utilizing hi-c and related technologies to easily analyze their datasets to search for TADs with differentially enriched contacts between hi-c experiments. It provides two different approaches to assessing significance of the change: both the empirical, permutation based test proposed by [Dixon *et al.*, 2015] and the parametric based on the Poisson-binomial distribution. Since the tool also allows for visualisation of results it should be useful to many scientists.

## Acknowledgements

We would like to thank Minna U. Kaikkonen and Henri Niskanen from The University of East Finland for providing us with their unpublished Hi-C data to test our method.

## Funding

This work was supported by the Polish National Center for Research and Development grant No. ERA-NET-NEURON/10/2013.

